# Dual effects of presynaptic membrane mimetics on α-synuclein amyloid aggregation

**DOI:** 10.1101/2021.07.13.452016

**Authors:** Yuxi Lin, Dai Ito, Je Min Yoo, Mi Hee Lim, Woo Kyung Yu, Yasushi Kawata, Young-Ho Lee

## Abstract

Aggregation of intrinsically disordered α-synuclein (αSN) under various conditions is closely related to synucleinopathies. Although various biological membranes have shown to alter the structure and aggregation propensity of αSN, a thorough understanding of the molecular and mechanical mechanism of amyloidogenesis in membranes remains unanswered. Herein, we examined the structural changes, binding properties, and amyloidogenicity of three variations of αSN mutants under two types of liposomes, 1,2-dioleoyl-sn-glycero-3-phosphocholine (DOPC) and presynaptic vesicle mimetic (Mimic) membranes. While neutrally charged DOPC membranes elicited marginal changes in the structure and amyloid fibrillation of αSNs, negatively charged Mimic membranes induced dramatic helical folding and biphasic amyloid generation. At low concentration of Mimic membranes, the amyloid fibrillation of αSNs was promoted in a dose-dependent manner. However, further increases in the concentration constrained the fibrillation process. These results suggest the dual effect of Mimic membranes on regulating the amyloidogenesis of αSN, which is rationalized by the amyloidogenic structure of αSN and condensation-dilution of local αSN concentration. Finally, we propose physicochemical properties of αSN and membrane surfaces, and their propensity to drive electrostatic interactions as decisive factors of amyloidogenesis.

## Introduction

α-Synuclein (αSN), an intrinsically disordered protein consisting of 140 amino acids is abundantly expressed in the brain. Although the exact function of αSN remains unclear, recent studies suggest that it plays an important role in modulating the neurotransmitter release (Burre, 2015) and protecting nerve terminals (Chandra et al., 2005). However, when exposed to stress conditions such as high levels of reactive oxygen species, soluble αSN monomers aggregates into insoluble amyloid fibrils with highly-ordered cross-β structures (Scudamore and Ciossek, 2018). Other forms of aggregates including oligomers are also observed as an intermediate in the process of amyloid fibrillation or as a dead-end product. The abnormal *in vivo* accumulation of αSN is the pathological hallmark of synucleinopathies including Parkinson’s disease (PD), dementia with Lewy bodies, and multiple system atrophy (MSA).

The self-assembly of αSN into amyloid fibrils is characterized by two sequential steps: slow nucleation followed by rapid elongation. It is generally accepted that physicochemical and biological factors exert significant impacts on the aggregation kinetics and pathways of αSN. Namely, previous studies indicate that lagged amyloid fibril formation under physiological conditions can be accelerated by increasing temperature to 57 °C or decreasing pH to 2.0 (Uversky et al., 2001). The presence of preformed amyloid seeds of lysozyme and insulin also promotes amyloidogenesis of αSN (Yagi et al., 2005). On the other hand, graphene quantum dots (GQDs), a promising carbon-based nanomaterial in biomedicine, prevent the aggregation of αSN monomers to amyloids (Kim and Yoo et al., 2018). In addition to αSN, amyloid beta (Aβ) and tau also display context-dependent aggregation behaviors (Lin et al., 2019;Gee and Lin et al., 2020).

Despite highlighted expression patterns in presynaptic terminals, αSN is widely distributed in the intracellular environment and interacts with various subcellular components. Among them, lipid membranes have been increasingly accentuated due to their critical impact on the structure and aggregation propensity of αSN. Upon binding to lipid membranes, the amphipathic N-terminal region (NTR) (residues 1 - ~60) and the hydrophobic non-amyloid β component (NAC) domain (residues ~60 - ~100) adopt α-helical structures (Dikiy and Eliezer, 2012). Based on the atomic level analysis, they manifest in two different helical conformations, broken helix and single elongated helix (Pfefferkorn et al., 2012). The distinct structures of αSN can be attributed to distinctive intermolecular interactions with membranes, which, in turn, dictate the amyloidogenicity of αSN. Along the same lines, the ratio of lipids to proteins (Galvagnion et al., 2015) and other chemical properties of membranes including the charge of head groups (Galvagnion et al., 2016) collectively influence the structure and amyloidogenesis of αSN. Moreover, our recent studies revealed that helical conformations in the initial structures of αSN in membranes is key to amyloid formation (Terakawa, Lee, and Kinoshita et al., 2018a).

Mutations in amyloid precursors are also crucial for regulating amyloidogenicity. For αSN, A53T and H50Q are the representative familial mutants associated with the early onset of PD, which manifest distinct aggregation behaviors and kinetics (Flagmeier et al., 2016;Meade et al., 2019). Truncated forms of αSN are also observed in Lewy bodies in cells where a truncation at the C-terminal leads to accelerated amyloid formation (Li et al., 2005;Izawa et al., 2012;Sorrentino et al., 2018). Other reports investigate the function of highly acidic C-terminal regions of αSN in membrane binding and subsequent amyloid formation. Even upon binding to membranes, the C-terminal domain remains disordered by making only weak and transient contacts with membrane surfaces (Fusco et al., 2014). Interestingly, the removal of the C-terminal regions remarkably reshapes the kinetic factors of the aggregation propensity under membrane environments. Although recent advances in characterization techniques have promoted our understanding of the effects of biological membranes on the aggregation of αSN, much remains uncertain about the molecular and mechanical mechanisms of amyloidogenesis of αSN in membranes.

Herein, we investigated mainly the impacts of presynaptic vesicle-mimicking model (Mimic) membranes on the amyloid fibrillation of αSN. Collective results from the structural change, membrane binding, and amyloid fibrillation of three αSN variants demonstrated that negatively charged Mimic membranes induce biphasic modulation of the amyloidogenicity of αSN. To explain this dual effect, *i.e*., promotion and inhibition, we propose two mechanisms based on the amyloidogenic structure of αSN and the condensation-dilution of local αSN concentration in membranes. Taken together, this study establishes a general mechanistic perspective on the amyloid fibrillation of αSN in membranes and thereby contributes to the rational design of candidates against its deleterious aggregation.

## Materials and Methods

### Materials

The full-length human αSN (αSN_WT_) and three variations of αSN mutants: 1) C-terminal 11-residue truncation (αSN_129_); 2) charge neutralization of negatively-charged residues between positions 130 and 140 to asparagine residues (αSN_130CF_); 3) mutation of the 53^rd^ residue from alanine to threonine (αSN_A53T_), were expressed in *E. coli* BL21 (DE3), and purified as previously described (Izawa et al., 2012). Phospholipids, DOPC, 1,2-dioleoyl-*sn*-glycero-3-phosphoethanolamine (DOPE), and 1,2-dioleoyl-*sn*-glycero-3-phospho-l-serine (DOPS) were obtained from Avanti Polar Lipids Inc. (Alabaster, USA) (Fig. S1). Thioflavin T (ThT) was purchased from Wako Pure Chemical Industries, Ltd (Osaka, Japan). All other reagents were obtained from Nacalai Tesque (Kyoto, Japan).

### Vesicle preparation

Small unilamellar vesicles (SUVs) containing DOPC or DOPC:DOPE:DOPS at a ratio of 2:5:3 were prepared as mimicking presynaptic vesicles according to the previous literature (Terakawa, Lee, and Kinoshita et al., 2018a). Briefly, lipids were dissolved in chloroform, and mixed in glass tubes at the desired compositions. The resulting solution was dried under a nitrogen stream, followed by vacuum drying to ensure the removal of residual organic solvents. To rehydrate the resultant lipid film, a solution of 20 mM sodium phosphate buffer (pH 7.4) containing 100 mM NaCl was added with vortex mixing. After 10 freeze-thaw cycles, lipid suspensions were sonicated for 10 mins on ice to obtain a homogeneous SUVs solution.

### ThT fluorescence assay

αSNs were dissolved in 20 mM sodium phosphate buffer (pH 7.4) containing 100 mM NaCl to prepare a stock concentration of 200 μM. Protein concentrations were determined using the UV-absorbance at 280 nm with molar extinction coefficients of 2,980 M^−1^∙cm^−1^ for αSN_129_, and 5,960 M^−1^∙cm^−1^ for αSN_WT_, αSN_130CF_, and αSN_A53T_. The following experimental conditions were used to investigate αSNs amyloid formation at 37 °C: 50 μM αSNs, 20 mM sodium phosphate buffer (pH 7.4), 100 mM NaCl, 5 μM ThT, and Mimic and DOPC model membranes at various concentrations of lipids. Sample solutions (200 μl) were applied in triplicate to each well of the 96-well microplate (Greiner-Bio-One, Tokyo, Japan), and sealed with a film (PowerSeal CRISTAl VIEW, Greiner-Bio-One, Tokyo, Japan). The microplate, placed on a water bath-type ultrasonic transmitter (Elestein SP070-PG-M, Elekon Sci. Inc., Chiba, Japan), was subjected to cycles of ultrasonication for 1 min at 9-min intervals. The fluorescence intensity of ThT was hourly recorded on an SH-9000 microplate reader (Corona Electric Co., Ibaraki, Japan) with excitation and emission wavelengths of 450 and 485 nm, respectively.

After the data acquisition, kinetic analyses of αSNs amyloid formation were carried out using the following equation:

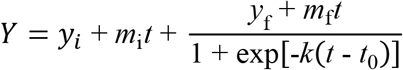

where *y*_i_ + *m*_i_*t* and *y*_f_ + *m*_f_*t* are the initial and final baselines, respectively. *t*_0_ is the half-time at which ThT fluorescence reaches 50% of the maximum amplitude. *k* represents the elongation rate constant. The lag time was obtained based on the following relationship: lag time = *t*_0_ - 2(1/*k*).

### Isothermal titration calorimetry

Isothermal titration calorimetry (ITC) experiments were performed with an ITC_200_ instrument (Malvern Panalytical, UK) at 25 °C. The concentrations of αSN in the ITC syringe and lipids of Mimic membranes in the ITC cell were 400 μM and 2 mM, respectively. αSNs were dissolved in 20 mM sodium phosphate buffer (pH 7.4) containing 100 mM NaCl. The reference power was set to 10 μcal∙sec^−1^, and the initial delay was 300 secs. Titration experiments consisted of 20 injections spaced at intervals of 300 secs. The injection volume was 0.4 μl for the first injection and 2 μl for the residual injections. The stirring speed was 1,000 rpm. Data were analyzed with a one-set of sites binding model using the MicroCal PEAQ-ITC Analysis Software (Malvern Panalytical, UK).

## Results

### Structural characterization of αSN mutants under membrane environments

Far-UV circular dichroism (CD) spectroscopy elucidates the effects of Mimic and DOPC membranes on the initial structures of three different αSN variants – αSN_129_, αSN_130CF_, and αSN_A53T_ (Fig. 1A, E, and I). In the absence of membranes, αSN_129_ exhibited a single negative band at ~200 nm without any noticeable band in the region between 210 and 230 nm, indicating that the secondary structures are predominantly disordered. On the other hand, increasing the concentration of Mimic lipids from 0 to 5 mM induced helix-rich conformations as characterized by the two negative bands at ~208 and ~222 nm (Fig. 1A, left). Further secondary structure analysis showed consistent results with increased helical structures and decreased β- and random-coil structures as a function of Mimic lipids concentration (Fig. S2).

**Figure 1.**
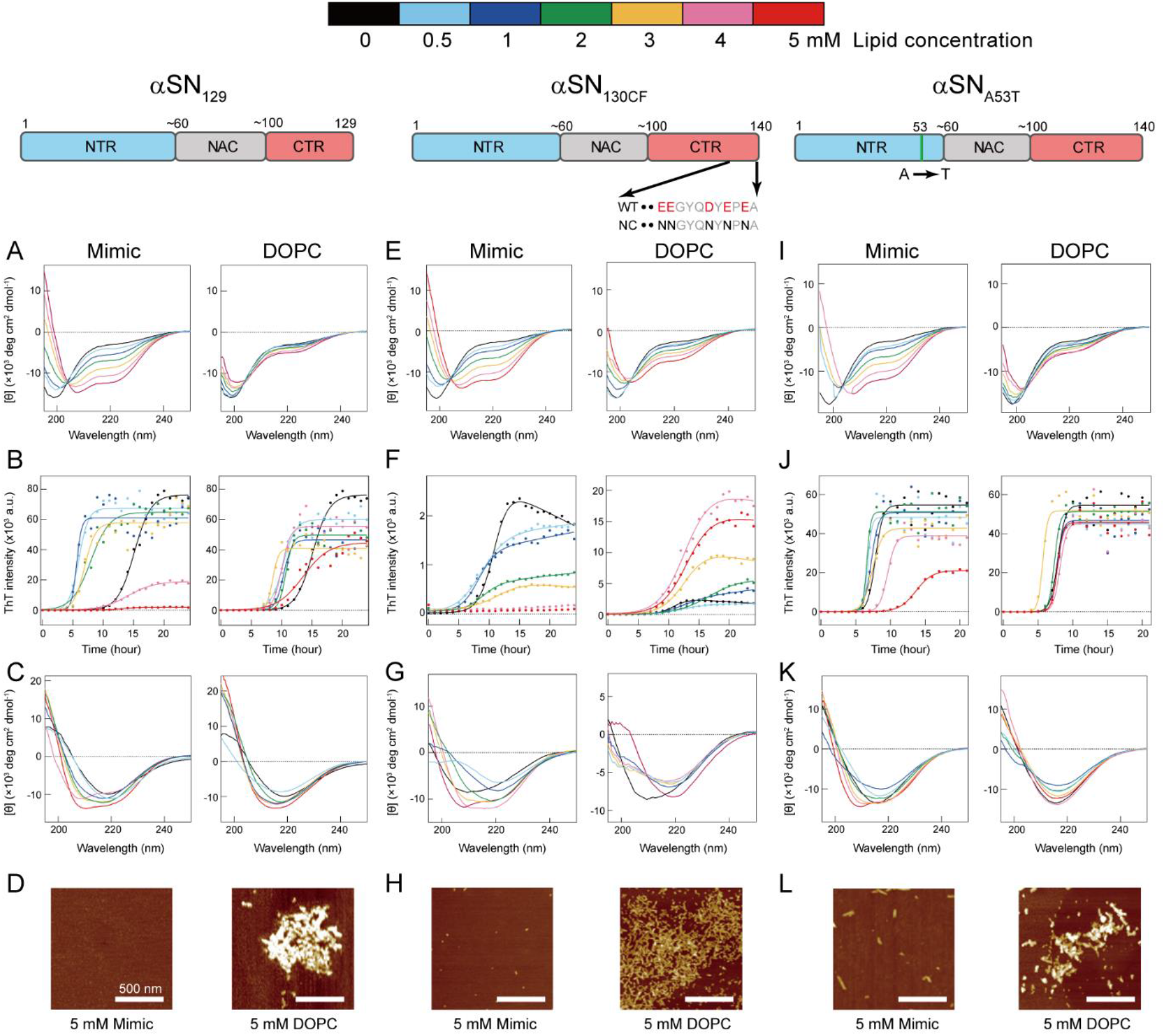
Effects of model membranes on the structure and amyloid formation of αSNs. (**A-L**) Conformational transitions and fibrillation kinetics of αSN_129_ (**A**-**D**), αSN_130CF_ (**E**-**H**), and αSN_A53T_ (**I-L**) in the absence and presence of Mimic (left) and DOPC membranes (right). Far-UV CD spectra of αSN_129_ (**A** and **C**), αSN_130CF_ (**E** and **G**), and αSN_A53T_ (**I** and **K**) before (**A**, **E**, and **I**) and after (**C**, **G**, and **K**) incubation were acquired. (**B**, **F**, and **J**) Fibrillation kinetics of αSN_129_ (**B**), αSN_130CF_ (**F**), and αSN_A53T_ (**J**) were monitored by the ThT fluorescence assay. Raw data averaged from three independent experiments are shown as closed circles. Solid lines represent the fit curves. Schematic representations of αSN_129_, αSN_130CF_, and αSN_A53T_ are displayed above the corresponding data. The *N*-terminal region (NTR), the non-amyloid β component (NAC) region, and the *C*-terminal region (CTR) are colored in blue, grey, and red, respectively. Various concentrations of lipids in Mimic and DOPC membranes are guided by distinct colors: black (0 mM), light blue (0.5 mM), blue (1 mM), green (2 mM), yellow (3 mM), pink (4 mM), and red (5 mM). (**D**, **H**, and **L**) AFM images were taken for the samples of αSN_129_ (**D**), αSN_130CF_ (**H**), and αSN_A53T_ (**L**) incubated with 5 mM of Mimic (left) or DOPC (right) lipids. The white scale bars indicate 500 nm.

In contrast to Mimic membranes, DOPC membranes caused negligible intensity magnifications in the negative peaks of CD spectra. Even after increasing the concentration of DOPC lipids to 5 mM, a minor structural alteration of αSN_129_ upon binding was still elicited (Fig. 1A, right). Similar structural reconfigurations to those of αSN_129_ were also observed for αSN_130CF_ and αSN_A53T_ in the presence of Mimic and DOPC membranes (Fig. 1E and I). These results indicate that Mimic membranes are more effective in generating helical structures of αSNs, which corroborate our previous findings with αSN_WT_ (Terakawa, Lee, and Kinoshita et al., 2018a).

### Amyloid formation of αSN mutants under membrane environments

ThT fluorescence assay examines the aggregation behaviors of the three αSN mutants in the absence and presence of the test membranes. In the absence of the membranes, the fluorescence intensities of αSN_129_, αSN_130CF_, and αSN_A53T_ increased after a lag phase at ~12-, ~8-, and ~7-hour post-incubation, and reached a plateau at ~20, ~13, and ~10 hours after incubation, respectively (Fig. 1B, F, and J). These typical sigmoidal growth curves indicate nucleation-dependent amyloid formation, which was also observed for the amyloid fibrillation of αSN_WT_ in the previous result (Terakawa, Lee, and Kinoshita et al., 2018a). In addition, the post-incubation far-UV CD spectra exhibited a single negative band near 218 nm, representing the cross-β structure of amyloid fibrils (Fig. 1C, G, and K). Collectively, the amyloid generation of all three αSN variants was verified in the absence of membranes.

The presence of Mimic membranes led to more dynamic alterations in the amyloidogenicity of αSN_129_. ThT fluorescence analysis revealed two distinct effects of Mimic membranes on fibrillation kinetics: 1) accelerated amyloid formation at lower concentrations of Mimic lipids (0.5 - 4 mM) with a shorter lag time and larger elongation rate constant; 2) constrained amyloid generation at higher concentrations (5 mM) with a more extended lag time and lower elongation rate constant (Figs. 1B and 2A-C, left). Such results correspond to the previous finding on lipid concentration-dependent amyloidogenesis of αSN_WT_ (Terakawa, Lee, and Kinoshita et al., 2018a). In accordance with the ThT assay results, the far-UV CD spectra at 0.5 - 3 mM and 4 - 5 mM of Mimic lipids revealed, respectively, amyloid fibrils with β-structures and monomers with predominant helical conformations (Fig. 1C, left). The atomic force microscopy (AFM) image at 5 mM of Mimic lipids further confirmed their inhibitory effects against amyloidogenesis (Fig. 1D, left).

**Figure 2.**
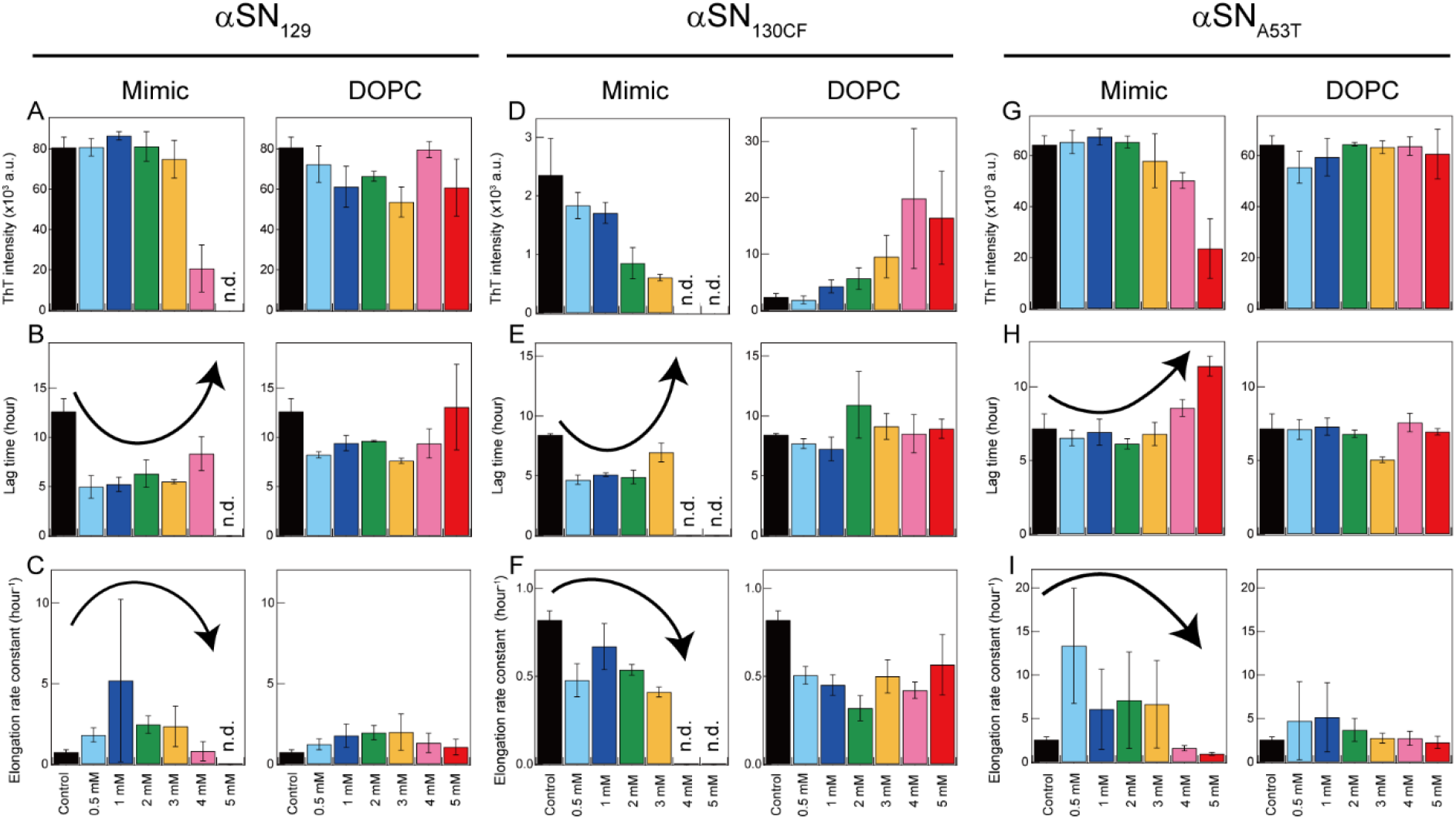
Kinetic analysis of amyloid formation of aSNs in model membranes. (**A**-**I**) Maximum ThT fluorescence intensities (**A**, **D**, and **G**), lag times (**B**, **E**, and **H**), and elongation rate constants (**C**, **F**, and **I**) of amyloidogenesis of αSN_129_ (**A**-**C**), αSN_130CF_ (**D**-**F**), and αSN_A53T_ (**G**-**I**) in the absence and presence of Mimic (left) or DOPC membranes (right). Average values calculated from three separate samples of a single set are shown with error bars reporting the standard deviation. “n.d.” denotes the concentration of lipids at which no significant increase in the ThT fluorescence intensity throughout the incubation period was observed. Various concentrations of lipids in Mimic and DOPC membranes are guided by distinct colors: black (0 mM), light blue (0.5 mM), blue (1 mM), green (2 mM), yellow (3 mM), pink (4 mM), and red (5 mM). The arrows indicate a trend of changes in the lag time (**B**, **E**, and **H**) and elongation rate constant (**C**, **F**, and **I**).

In contrast to Mimic membranes, DOPC membranes exhibited minimal effects on the amyloid fibrillation of αSN_129_. As shown in Figure 1B (right), similar nucleation-dependent sigmoidal increases in the ThT intensity were observed at all DOPC lipid concentrations (0.5 – 5 mM). Further kinetic analyses verified that DOPC lipids at the concentration range between 0.5 and 5 mM slightly promoted αSN_129_ amyloid formation by affecting the lag time (Fig. 2B, right). The far-UV CD spectra of αSN_129_ at all concentrations of DOPC lipids showed the formation of cross-β-structured amyloid fibrils after incubation (Fig. 1C, right). It was further verified by AFM analysis at 5 mM of DOPC lipids, which exhibited clustered amyloid fibrils (Fig. 1D, right). Similar minimal effects of DOPC membranes were also revealed for αSN_WT_ aggregation in the previous literature (Terakawa, Lee, and Kinoshita et al., 2018a).

Next, we investigated the effects of Mimic and DOPC membranes on αSN_130CF_ amyloid formation. The addition of 0.5 - 2 mM of Mimic lipids remarkably accelerated amyloid fibrillation by shortening the lag time from ~8 to ~4 hours (Figs. 1F and 2E, left). However, increased lipid concentrations (4 – 5 mM) impeded the fibrillation of αSN_130CF_, leading to no increase in the ThT intensity throughout incubation. Consistent with ThT results, far-UV CD spectra at upper range lipid concentrations showed predominant helical structures (Fig. 1G, left) without detectable fibrillar aggregates at 5 mM of Mimic lipids (Fig. 1H, left). On the other hand, the presence of DOPC membranes did not yield noticeable changes on the amyloid formation of αSN_130CF_. At all concentrations of DOPC lipids, ThT intensities increased after a lag time of ~8 - ~11 hours (Figs. 1F and 2E, right). Although the maximum ThT intensities at high concentrations of DOPC lipids were greater than those at low and middle concentrations (Fig. 2D, right), such discrepancy can be attributed to the polymorphic nature of amyloid fibrils. Along the same lines, cross-β structures were detected at all DOPC concentrations after incubation (Fig. 1G, right), with evident fibrillar aggregates formation at 5 mM of DOPC (Fig. 1H, right).

Analogous to the findings for αSN_129_ and αSN_130CF_, Mimic membranes accelerated and inhibited the amyloidogenesis of αSN_A53T_ in a concentration-dependent manner. As the concentration of Mimic lipids increased from 0 to 5 mM, the lag time initially decreased and then increased, with the elongation rate constant showing the opposite tendency (Figs. 1J, 2H and I, left). While some fibrillar fragments were observed at 5 mM of Mimic lipids (Fig. 1L, left), the majority of αSN_A53T_ existed as helical monomers (Figs. 1K and 2G, left). On the contrary, almost no effect of DOPC membranes on the amyloid formation of αSN_A53T_ was detected. Kinetic analyses of the ThT data revealed that the lag time and elongation rate constant of αSN_A53T_ fibrillation remained steady throughout all lipid concentrations (Fig. 2H and I, right). In like manner, far-UV CD spectra indicated the existence of fibrillar aggregates with cross-β structures at all DOPC concentrations (Fig. 1K, right), which was supported by the representative AFM image in the presence of 5 mM of DOPC lipids (Fig. 1L, right).

### Calorimetry-based investigation of intermolecular interactions between Mimic membranes and αSNs

To obtain further insights into how Mimic membranes modulate the amyloidogenicity of αSN, we performed ITC analysis on αSN-membrane interactions. As shown in Figure 3A-D (upper), a series of titration of αSNs to Mimic membranes generated negative ITC peaks followed by gradual saturation. This suggests the presence of appreciable exothermal intermolecular interactions between αSNs and Mimic membranes. Following normalization of all ITC peaks, ITC thermograms were converted to binding isotherms (Fig. 3A-D, lower). The thermodynamic parameters obtained from fitting the binding isotherms are summarized in Figure 3E.

**Figure 3.**
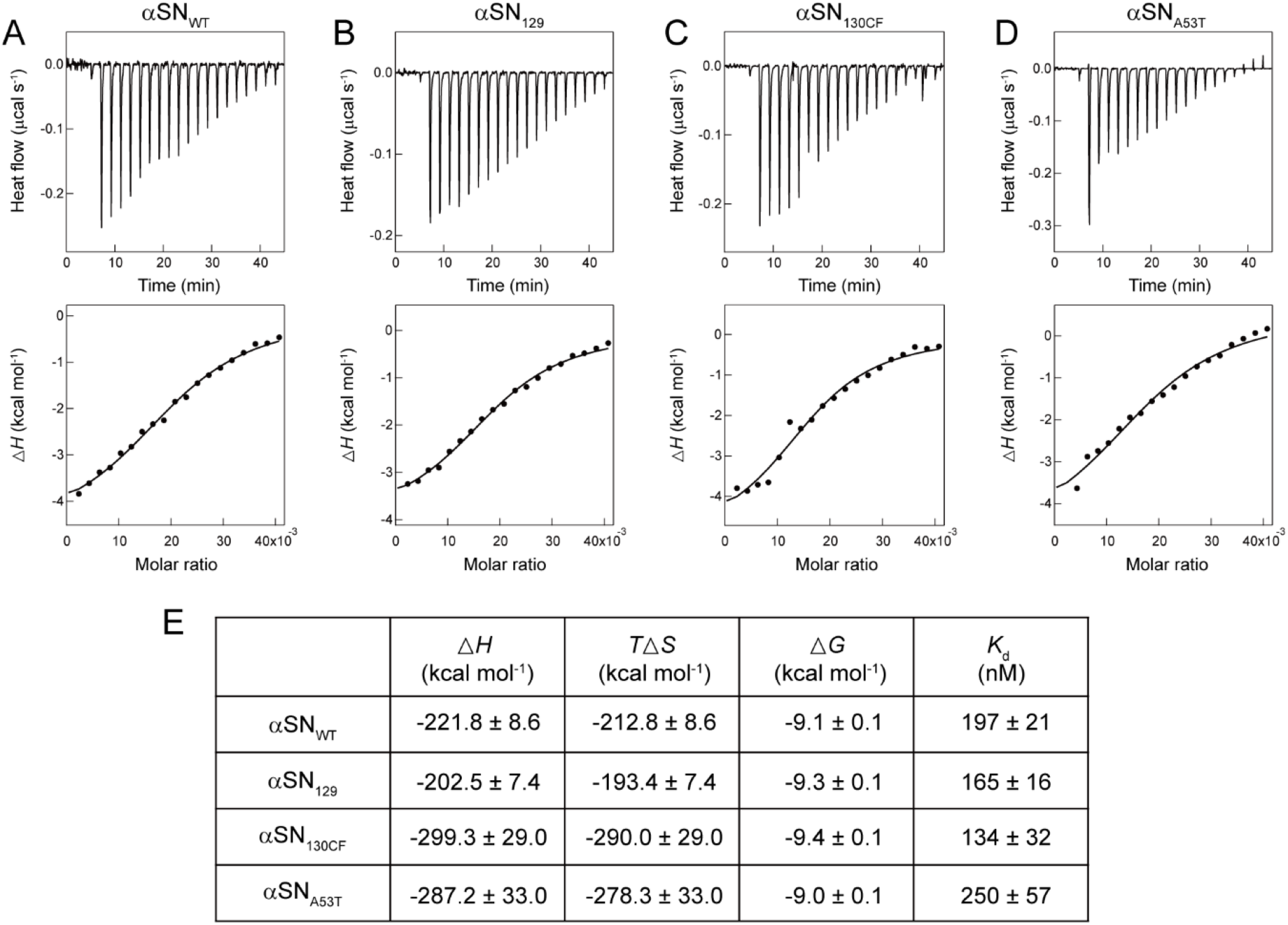
Calorimetry-based characterization of interactions between αSNs and Mimic membranes. (**A**-**E**) ITC thermograms (upper) and binding isotherms (lower) obtained by titrating αSN_WT_ (**A**), αSN_129_ (**B**), αSN_130CF_ (**C**), and αSN_A53T_ (**D**) to Mimic membranes are shown. Solid lines in binding isotherms indicate the fit curves based on a one-set of sites binding model. (**E**) Thermodynamic parameters for the binding of αSNs to Mimic membranes.

As expected from downward ITC peaks, thermodynamically favorable enthalpy changes (Δ*H* < 0), ranging from ~−200 to ~−300 kcal∙mol^−1^, ensued for all variants of αSN. A large negative value of Δ*H* may stem from the membrane binding of αSNs being steered by attractive electrostatic interactions and concomitant helical folding. On one hand, unfavorable negative entropy change (*T*Δ*S* < 0) were monitored from ~−190 to ~−290 kcal∙mol^−1^. Nevertheless, the loss of conformational and translational entropies owing to the membrane-induced helical folding of αSNs and the restricted lipid diffusion were compensated by large negative Δ*H* resulting in thermodynamic stabilization. The outcomes demonstrate that αSNs-Mimic membrane interactions are purely driven by the enthalpy change. Similar trends entailed for the interactions between αSN_WT_ and model membranes which consisted of phosphatidylserine and gangliosidosis-1 (Nuscher et al., 2004;Bartels et al., 2014).

ITC analyses provided distinct dissociation constant (*K*_d_) for all binding systems which are within a similar range. The changes in the Gibbs free energy (Δ*G*) showed negative values ranging from −9.0 to −9.4 kcal∙mol^−1^, which indicate spontaneous interactions of all αSN variants with Mimic membranes. It should be noted that the binding affinity decreased in the order of αSN_130CF_, αSN_129_, αSN_WT_, and αSN_A53T_. Altogether, the findings from the ITC study suggest that the negative charges of the C-terminal region play a pivotal role in the thermodynamic adjustment of αSN upon binding to Mimic membranes.

## Discussion

We investigated the impacts of lipid membranes on the amyloid formation of three variations of αSNs (αSN_129_, αSN_130CF_, and αSN_A53T_) with different charge states and lengths as functions of lipid component and concentration. Based on the structural, kinetic, and thermodynamic characterizations, the molecular and mechanical mechanisms of membrane-assisted acceleration and inhibition of amyloid generation were elucidated (Figs 1, 2, and S1). While neutrally charged DOPC membranes showed insignificant effects on the structure and amyloidogenicity of αSNs, negatively charged Mimic membranes induced dramatic helical transitions with the dual effects of promoting and impeding amyloid aggregation depending on the membrane concentration. At low concentrations of Mimic lipids, the fibrillation of αSNs was accelerated, whereas high lipid concentrations abrogated the process. Similar dual effects on the amyloidogenicity of αSN_WT_ were reported for other membranes with negatively charged lipids such as DOPS and DMPS (Galvagnion et al., 2015;Galvagnion et al., 2016), as well as SDS (Giehm et al., 2010). Thus, these findings strongly implicate lipid concentration-dependent acceleration and inhibition of αSN amyloidogenesis under membrane-binding conditions with a net negative charge of the head groups.

To rationalize the dual effects of negatively charged Mimic membranes, we conceive two possible mechanisms based on the initial structure of αSN in membranes (the amyloidogenic structure) and the intermolecular affinity of αSN for membranes (condensation-dilution). The initial structure model elucidates the dual effect based on distinct structures of αSNs at varying lipid concentrations (Fig. 4A). In the absence of Mimic lipids, largely disordered αSNs slowly self-assemble into amyloid fibrils with cross-β structures. The addition of Mimic lipids at low concentrations triggers structural alteration from random coils to partial helical structures (Fig. 4A, upper). Partial helical structures are inclined to interact with one another through helix-helix interactions, thereby facilitating nucleation for amyloidogenesis (Abedini and Raleigh, 2009;Lin et al., 2019).

**Figure 4.**
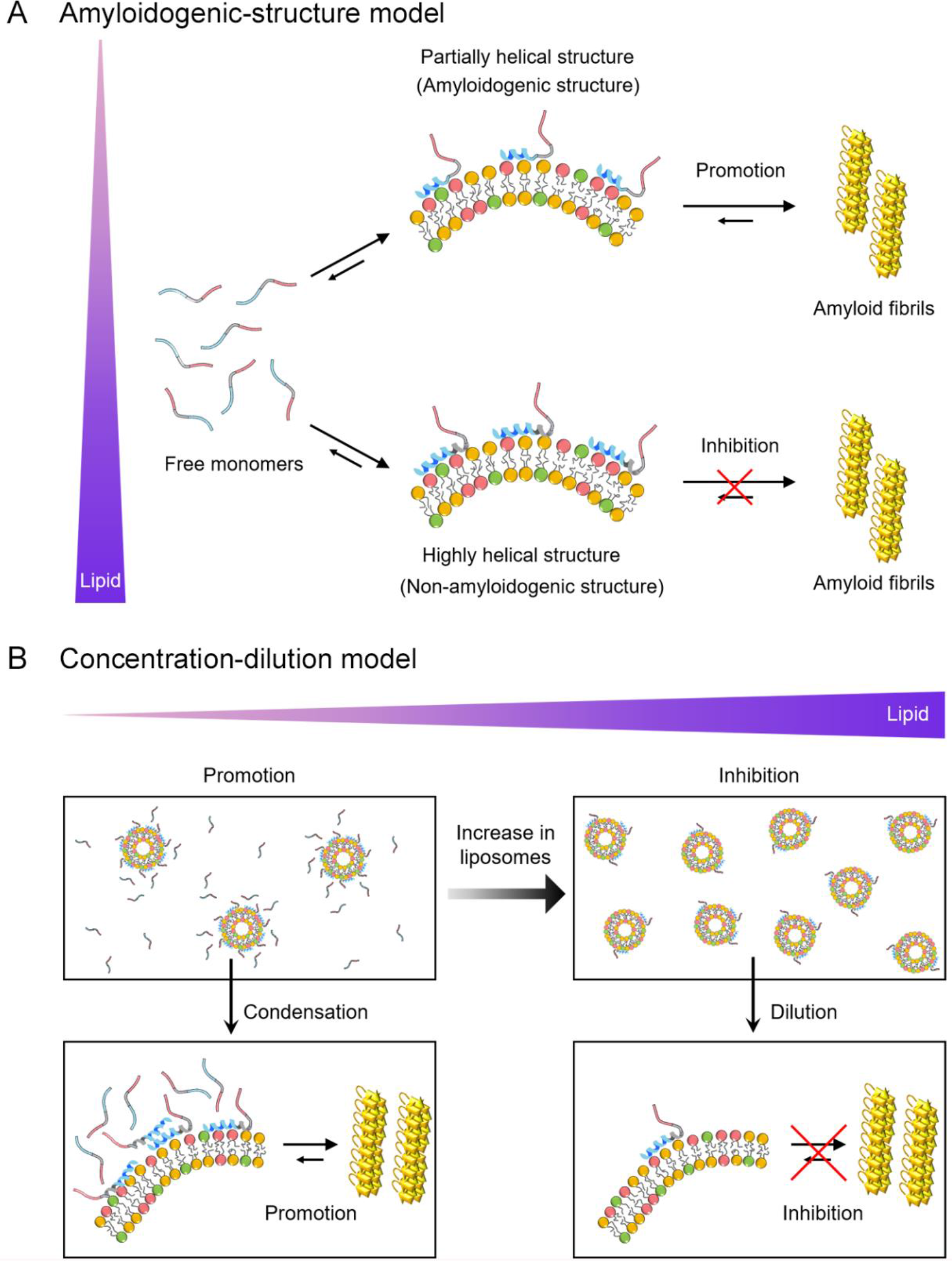
Schematic models for the dual effect of Mimic membranes on αSN amyloidogenesis. (**A** and **B**) Two models, the amyloidogenic structure model (**A**) and the condensation-dilution model (**B**) are schematically shown. Free monomers, partially- and highly-helical monomers in the membrane-bound forms, and amyloid fibrils are illustrated. The *N*-terminal region, the non-amyloid component region, and the *C*-terminal region of αSN are represented in blue, grey, and red, respectively. Increases in the concentration of lipids are indicated by the purple triangle.

Previous studies also suggested that partial helical structures are aggregation-prone and are the representative secondary structures of the key intermediates in the fibrillation pathway of αSN (Anderson et al., 2010;Ghosh et al., 2015), Aβ40 (Lin et al., 2019), hIAPP (Pannuzzo et al., 2013), and polyQ (Jayaraman et al., 2012), proposing an existence of amyloidogenic structure. Amyloidogenic structures have also been implicated in other folded proteins such as SH3 domain (Guijarro et al., 1998) and β2-microglobulin (Jahn et al., 2006). In contrast, at high concentrations of Mimic lipids, αSNs adopt prominent helical structures with an exceptionally low aggregation propensity (Fig. 4A, lower). These highly helical non-amyloidogenic structures were analogously observed in Aβ40, Aβ42, and αSNs at high concentrations of alcohols (e.g. 40% TFE and 50% HFIP) (Crescenzi et al., 2002;Anderson et al., 2010;Lin et al., 2019). Accordingly, αSNs in bulk aqueous solution would take time to form a nucleus with an amyloidogenic structure in a conformational ensemble.

Condensation-dilution model explains the mechanism of the dual effect on the basis of the thermodynamic binding affinity (Fig. 4B). At low lipid concentrations, αSN binds multiply with Mimic membranes, which leads to increased local concentrations of αSN (Fig. 4B, left). Thus, concentrated αSN will be sufficient to facilitate nucleation for amyloid fibrillation. The growth process can be expedited by elongation with the addition of neighboring monomers around fibril seeds. However, at high lipid concentrations, αSNs will be spread across discrete liposomes and their membranes, leading to diluted local concentrations of αSN. As a result, the amount of αSNs in each liposome and bulk water decrease significantly. This, in turn, interferes with efficient nucleation and elongation, causing the prevention of amyloid formation (Fig. 4B, right). In addition, the condensation-dilution model also illustrates the aggregation of Aβ at the various concentration of cationic polystyrene nanoparticles (Cabaleiro-Lago et al., 2010).

Biological membranes have shown their capability to modulate folding, aggregation, and the function of αSN (O’Leary and Lee, 2019). Binding affinity of αSN for membranes is influenced not only by the properties of lipid bilayers such as the net charge and curvature, but also by mutations and post-translational modifications including phosphorylation (Kuwahara et al., 2012) and N-terminal acetylation (Runfola et al., 2020). In the current study, we revealed that the binding affinity of αSNs to Mimic membranes decreased in the order αSN_130CF_, αSN_129_, αSN_WT_, and αSN_A53T_. This indicates that the removal of negatively charged residues between positions 130 and 140 increases the membrane binding affinity, whereas repulsive electrostatic interactions between negatively charged C-terminal domain of αSN and Mimic membranes decrease the intermolecular affinity. Considering that the large energy gain for αSN upon membrane binding is derived from electrostatic interactions between the positively charged NTR of αSN and negatively charged membranes, electrostatic forces are fundamental for αSN-membrane interactions. Increased affinity for membranes with an additional positive charge in the NTR of E46K further supports the importance of electrostatic contributions (Stockl et al., 2008). In addition, we speculate that a point mutation in the NTR like αSN_A53T_ might impair favorable electrostatic interactions with membranes, which attenuates the overall affinity.

In line with present results, a recent study reported that the addition of calcium ions significantly increases αSN’s propensity to interact with negatively charged membranes by reducing repulsive electrostatic interactions of negatively charged C-terminal regions with membranes (Lautenschlager et al., 2018). Along the same lines, the higher affinity of αSN_130CF_ can be attributed to possible contacts of neutralized 10 residues with membranes via non-polar interactions. It should be also noted that the minimal concentration of Mimic lipids for blocking fibrillation (αSN_130CF_: 4 mM; αSN_129_ ≈ αSN_WT_: 5 mM; αSN_A53T_: > 5 mM) mostly followed the reverse order of the binding affinity, which further supports the condensation-dilution model. Overall, the relative molar ratio of αSN to the lipid concentration is a decisive parameter of amyloid generation in presynaptic vesicles.

Phase diagrams are highly valuable for a comprehensive understanding biological and pathogenic phase transitions including protein aggregation (Lin and Lee et al., 2014;Lin et al., 2016;Terakawa, Lee, and Kinoshita et al., 2018a;Terakawa et al., 2018b;Lin et al., 2019;Gee and Lin et al., 2020;Ivanova et al., 2021). To illustrate membrane-induced amyloidogenesis of αSNs, we constructed conceptual phase diagrams of αSN_130CF_, αSN_129_, and αSN_A53T_ depending on the concentrations of αSN and Mimic lipids (Fig. S3). Each αSN displays soluble-to-insoluble phase transition following thermodynamic equilibration. Displaying amyloid-forming regions of αSN_130CF_, αSN_129_, and αSN_A53T_ at 50 μM respectively at 0 - 4, 0 - 3, and 0 - 5 mM of Mimic lipids demonstrate the minimal concentration of Mimic lipids required to impede the fibrillation process. Further elevations in the lipid concentration beyond the amyloid-forming region may increase the solubility of αSNs, thereby preventing their aggregation.

Understanding of context-dependent kinetics and amyloidogenicity of αSN is essential for overcoming synucleinopathies with cytotoxic aggregation in cells. Depending on its cellular localization and neighboring components, the amyloid fibrillation of αSN will be both faster and slower in bulk solution than in biological membranes, including presynaptic vesicles, due to the dual effect. As the dual effect based on two possible models suggests, αSN amyloid fibrillation is subjected to acceleration or inhibition depending on the structural state of αSN and its relative affinity for membranes. In summation, the modulation of amyloidogenesis is governed by various conditions that regulate electrostatic interactions between αSN and membranes through a favorable enthalpic contribution.

## Author Contributions

Y.L. and Y.-H.L. conceived the presented idea. Y.L. and D.I. carried out the experiment. D.I., M.H.L. J.M.Y., W.K.Y., and Y.K. contributed to the interpretation of the results and edited the manuscript. Y.L. and Y.-H.L. wrote the manuscript with input from all authors. All authors contributed to the article and approved the submitted version.

## Acknowledgments

We thank Dr. Mayu. S. Terakawa (Kyoto Univ., Japan) and Prof. Yuji Goto (Osaka Univ., Japan) for their help to the current study. We thank Prof. Masahiro Shirakawa (Kyoto Univ., Japan) for contributing to ITC experiments. The authors acknowledge J.P. Hostetler (BIOGRAPHENE, USA) for assisting in manuscript completion and revision.

