## Supplementary Material for "Dual effects of presynaptic membrane mimetics on α-synuclein amyloid aggregation"

### Supplementary Information
### for
### Dual effects of presynaptic membrane mimetics on α-synuclein amyloid aggregation

Yuxi Lin[1,2*], Dai Ito[3], Je Min Yoo[4], Mi Hee Lim[5], Woo Kyung Yu[3,6], Yasushi Kawata[7], Young-Ho Lee[1,2,8,9,10*]

[1]Research Center of Bioconvergence Analysis, Korea Basic Science Institute (KBSI), Ochang, Chungbuk 28119, Republic of Korea

[2]Institute for Protein Research, Osaka University, Yamadaoka 3-2, Suita, Osaka 565-0871, Japan

[3]Department of Brain and Cognitive Science, Daegu Gyeongbuk Institute of Science and Technology (DGIST), Daegu, 42988, Republic of Korea

[4]BIOGRAPHENE, 555 W. 5th St., Los Angeles, California 90013, United States.

[5]Department of Chemistry, Korea Advanced Institute of Science and Technology (KAIST), Daejeon 34141, Republic of Korea

[6]Core Protein Resources Center, Daegu Gyeongbuk Institute of Science and Technology (DGIST), Daegu, 42988, Republic of Korea

[7]Department of Chemistry and Biotechnology, Graduate School of Engineering, Tottori University, Tottori 680-8550, Japan

[8]Bio-Analytical Science, University of Science and Technology (UST), Daejeon 34113, Republic of Korea

[9]Graduate School of Analytical Science and Technology (GRAST), Chungnam National University (CNU), Daejeon 34134, Republic of Korea

[10]Research Headquarters, Korea Brain Research Institute (KBRI), Daegu 41068, Republic of Korea

#### 1. Supplementary Materials and Methods

##### Atomic force microscopy

After incubating αSN monomers with Mimic and DOPC membranes at 5 mM lipids, sample drops of 50 μM αSNs were deposited on freshly cleaved mica plates. Following 1 min, the remaining solution was blown off with compressed air and further air-dry. Atomic force microscopy images were acquired using a Digital Instruments Nanoscope IIIa scanning microscope (Veeco, Santa. Barbara, CA) with a Si microcantilever.

##### Circular dichroism spectroscopy

Circular dichroism (CD) experiments were carried out on a JASCO J820 spectrophotometer (Tokyo, Japan) at 37 $^{o}$C. The far-UV CD spectra of αSNs in 20 mM sodium phosphate buffer (pH 7.4) containing 100 mM NaCl and various concentrations of lipids of DOPC and Mimic membranes were recorded using a quartz cuvette with a 0.1-mm path length. After subtracting the solvent background, CD signals were presented as the mean residue ellipticity (deg·cm$^{2}$· dmol$^{-1}$). The content of secondary structures was predicted using the BeStSel algorithm (Micsonai et al., 2015).

#### 2. Supplementary Figures

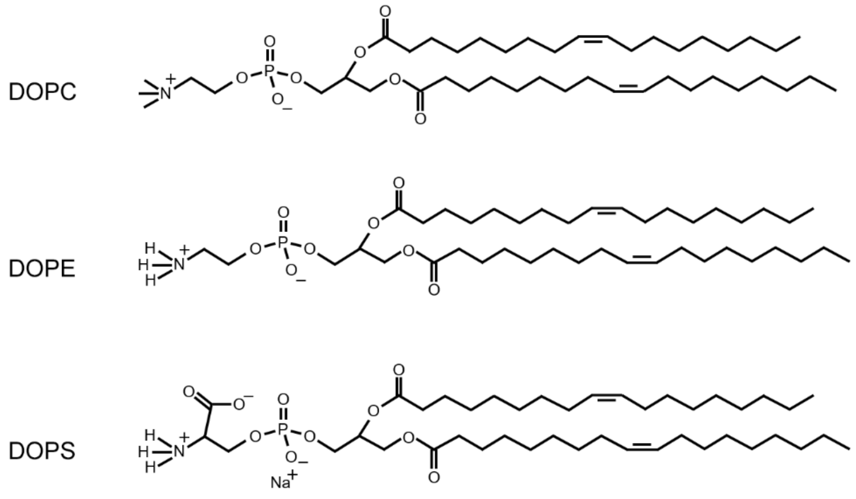


**Figure S1. Chemical structures of phospholipids.** Three types of phospholipids are shown: 1,2-dioleoyl-sn-glycero-3-phosphocholine (DOPC), 1,2-dioleoyl-*sn*-glycero-3-phosphoethanolamine (DOPE), and 1,2-dioleoyl-*sn*-glycero-3-phospho-l-serine (DOPS).

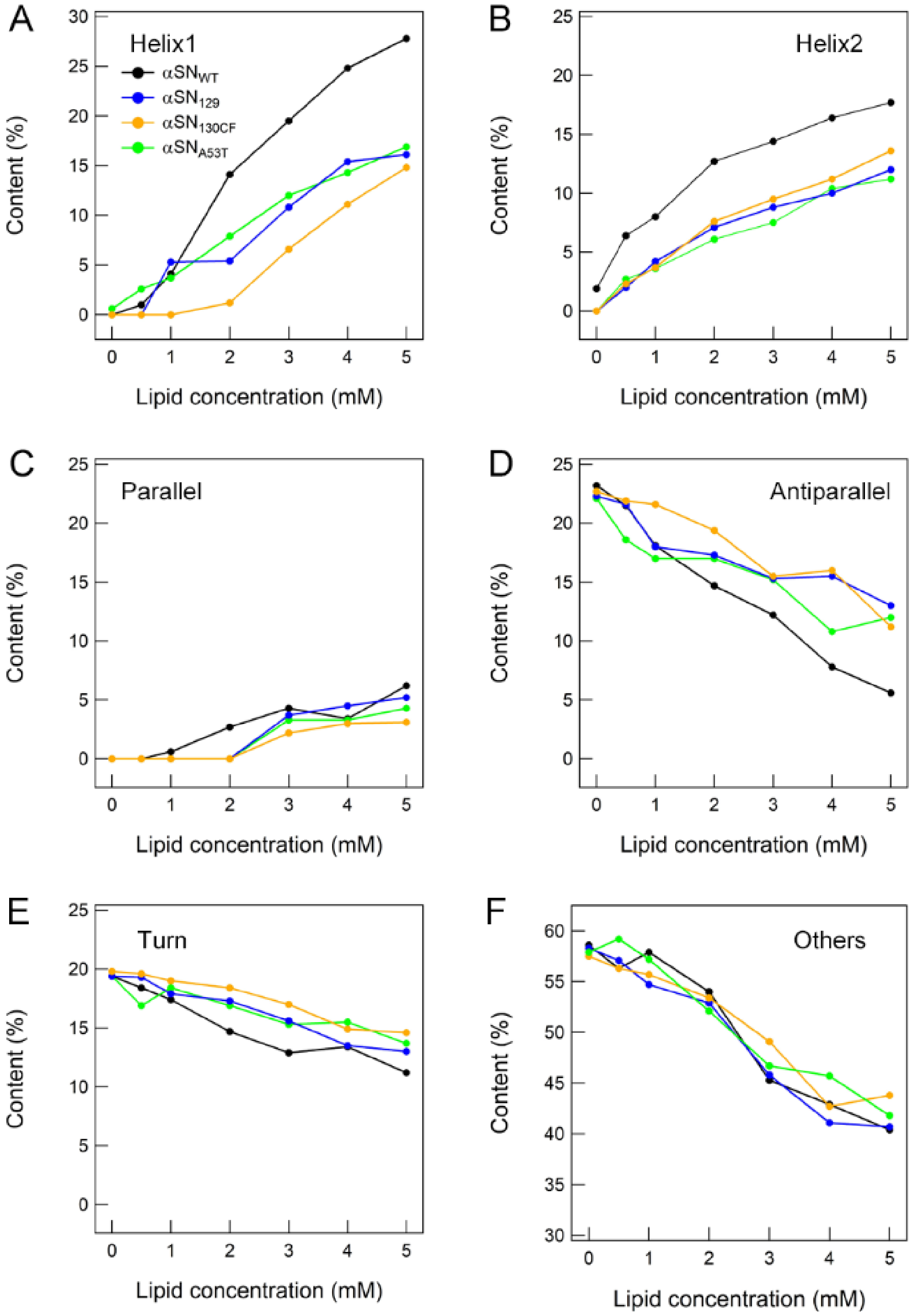
A
Helix1
αSN_WT
αSN_129
αSN_130CF
αSN_A53T
B
Helix2
C
Parallel
D
Antiparallel
E
Turn
F
Others
Content (%)
Lipid concentration (mM)

**Figure S2. Contents of the secondary structure of αSNs at the various concentrations of lipids of Mimic membranes**. (**A**-**F**) Contents of helix1 (**A**), helix2 (**B**), parallel β-sheet (**C**), antiparallel β-sheet (**D**), turn (**E**), and others (**F**) are plotted as a function of the concentration of Mimic lipids. Calculated contents of αSNs are displayed in distinct colors: $\alpha SN_{WT}$ (black), $\alpha SN_{129}$ (blue), $\alpha SN_{130CF}$ (yellow), and $\alpha SN_{A53T}$ (green). Results of $\alpha SN_{WT}$ were reproduced with modifications from our previous study (Terakawa, Lee, and Kinoshita et al., 2018).

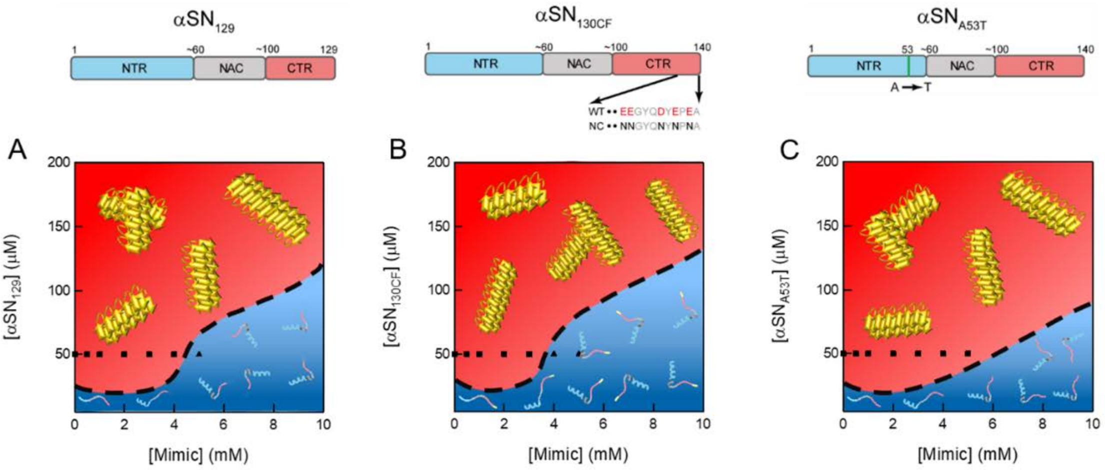


**Figure S3. Macroscopic phase diagrams of αSNs aggregation in Mimic membranes.** (**A**−**C**) Phase diagrams of amyloid formation of $\alpha SN_{129}$ (**A**), $\alpha SN_{130CF}$ (**B**), and $\alpha SN_{A53T}$ (**C**) in the presence of Mimic membranes. Colors and symbols represent the molecular species and colloidal states: soluble monomers (blue region and ▲) and mature amyloid fibrils (red region and ■). Symbols (■ and ▲) were plotted against the lipid concentration that was used for experiments. Cartoons for individual molecular species (largely-disordered monomer (bulk solution), highly-helical monomer (membrane-bound form), and amyloid fibril) are illustrated. Three distinct regions of αSN monomers are displayed in different colors: the N-terminal region (blue), the non-amyloid component region (grey), and the C-terminal region (red). Yellow color of $\alpha SN_{130CF}$ in the phase diagram indicates the region from E130 to A140 where negatively-charged residues are neutralized. Broken black lines at each phase diagram indicate conceptual solubility curves.
